# Measuring Phylogenetic Information Content Based on Clade Support

**DOI:** 10.64898/2026.11.13.766116

**Authors:** Johan A. A. Nylander

## Abstract

Phylogenetic information content is often understood as the degree to which data lend support to hypotheses about the monophyly of clades. This is most commonly expressed in terms of clade support, typically estimated by resampling techniques such as the bootstrap and the jackknife. Here, I discuss how to summarize this aspect of data quality. I show that raw bootstrap or jackknife proportions are unsuitable for this purpose and instead suggest a logarithmic transformation, which can be interpreted as translating the clade support values to the number of characters expected to give the same resampling proportion in parsimony analysis of a homoplasy-free data matrix. The distribution of the number of these “supporting characters” and descriptive statistics derived from this distribution, such as the mean (Mean Character Support, or MCS) and its standard deviation, provide powerful summaries of the quality of the underlying data set. I demonstrate for both empirical and simulated data sets that MCS accurately reflects information content, and that some popular alternative measures of data quality (the ensemble consistency index, the ensemble retention index, data decisiveness and the total support index) do not. Unlike most other measures of data quality, MCS can be applied to trees resulting from a range of alternative phylogenetic methods, including parsimony, distance and maximum likelihood approaches.

---

Phylogenetic systematists are often concerned with comparisons of the quality of different data sets (e.g. Friedlander et al., 1992, Graybeal, 1994). These sets could represent different sources of data (e.g. behavioral, morphological, and molecular data) or different partitions of the same data set (e.g. codon position in a protein-coding gene). Although different aspects of data quality might be of interest in these comparisons, the focus is commonly on the degree to which the data lend support to hypotheses about the monophyly of various clades. I will refer to this property as *phylogenetic information content*. It can be divided into two components: the degree of topological resolution, and the degree of support for the resolved clades.

Although it is not self-evident, most measures of data quality proposed thus far do not assess information content as defined above. For instance, it will be shown below that this is the case for the ensemble consistency index, *C* (Kluge and Farris, 1969) and the ensemble retention index, *R* (Archie, 1989a, Farris, 1989). Both these indices are based on the fit of individual characters to the shortest tree and measure average character quality, not information content per se. Furthermore, the values of *C* and *R* are dependent on both the number of characters and the number of terminals in a fashion that is not directly correlated with character quality, making comparisons between data sets difficult. Data sets with few characters tend to have larger *C* and *R* than data sets with many characters, even if average character quality is similar (Archie, 1989a, Klassen et al., 1991, Meier et al., 1991).

Other measurements of data quality focus on testing the presence of phylogenetic signal in the data and are based on evaluation of random trees or permuted or randomized data sets. The Homoplasy Slope Ratio (Meier et al., 1991), the *g*_1_ statistic (Hillis, 1991) and the Permutation Tail Probability (Archie, 1989b) are measures of this kind. There is considerable controversy surrounding the appropriateness of some of these tests (e.g., Källersjö et al., 1992; Swofford et al., 1996; Carpenter et al., 1998). For example, a randomly generated data matrix could be indicated to have significant hierarchical signal by the Permutation Tail Probability test, and yet have no supported groups (e.g., Carpenter et al., 1998:110). However, even if these test statistics were powerful indicators of phylogenetic signal (structure/hierarchy), it would be inappropriate to use them as estimators of information content.

Goloboff (1991:216) argued that “the worst data (the data which provide least information for a decision) are not the least hierarchical, but instead those for which no one scheme of relationships is better than any other.” He introduced the cladistic data decisiveness (DD) index to measure the amount of information in a data set that allows a choice between different trees, that is, a quantity that is potentially similar to phylogenetic information content as defined above. DD for a data set is calculated as *s̅* – *S*_MPT_ / *s̅* – *M*, where *s̅* is the mean length of all possible dichotomous trees, *S*_MPT_ the length of the most parsimonious tree, and *M* the minimum length of a tree without homoplasy (Goloboff, 1991). A data set without homoplasy will have a DD of 1 since *S*_MPT_ = *M*. DD is not a direct measure of homoplasy (Goloboff, 1991; De Laet and Smets, 1999), thus it potentially reflects data quality differently than indices such as *C* and *R*. This makes DD an interesting candidate measure for quality comparisons between data sets.

An common way of summarizing the results of a phylogenetic analysis is in terms of clade support values, that is, the degree to which the data lend support to hypotheses about the monophyly of various clades (taxon partitions). Clade support is typically assessed using a resampling scheme, either the bootstrap (Felsenstein, 1985) or the jackknife (Farris et al., 1996), or (for parsimony analyses) by Bremer support (decay index) (Bremer, 1988; 1994). There is an ongoing debate about the use of resampling techniques in phylogenetics (see Sanderson, 1995 for a review). Recent research has focused on the statistical properties of the techniques and the specific problems that arise when they are coupled with phylogenetic estimation. Such problems are, e.g. a general bias of the bootstrap (Zharkikh and Li, 1992; Felsenstein and Kishino, 1993; Hillis and Bull, 1993; Efron et al., 1996; Newton, 1996; Sanderson and Wojciechowski, 2000), shortcomings of the consensus technique used to summarize the results of the bootstrap or the jackknife (Wilkinson, 1996), and significant deviations depending on the heuristic search strategies employed (DeBry and Olmstead, 2000; Mort et al., 2000; Sanderson and Wojciechowski, 2000).

Despite the problems and assumptions associated with their estimation, it is clear that clade support values provide powerful and intuitive measures of relative strength for phylogenetic groupings. It is surprising, then, that there has been little discussion about measures summarizing phylogenetic information content based on clade support values. The only work I am aware of deals with Bremer support (Källersjö et al., 1992; Bremer, 1994), a measure that cannot be interpreted in probabilistic terms. Here I develop a measure of information content, the Mean Character Support (MCS), which is derived from logarithmic transformations of bootstrap- or jackknife-resampling proportions. I demonstrate for both empirical and simulated data sets that MCS is correlated with the degree to which data lend support to hypotheses about the monophyly of clades. I also show that some alternative indices, such as *C*, *R*, DD and a measure derived from Bremer support, do not measure this aspect of data quality. MCS is easily calculated, has a straightforward interpretation, and the underlying distribution from which it is calculated gives a comprehensive graphical summary of the information content of the underlying matrix.

## Example Data Sets

Three sets of data matrices will be used in illustrating the properties of data quality indices. The first is a set of molecular sequences for some *Drosophila* species (Baker and DeSalle, 1997). The subsets of this data matrix vary considerably in information content as is evidenced by the degree of resolution and clade support obtained when analyzing them separately. The total-evidence analysis is considerably richer in phylogenetic information than the smaller data sets. This is seen both in higher resolution and in much improved clade support in this analysis. However, this increase in information content is not picked up by all indices of data quality, as will be shown below.

Two simulated sets of data matrices were also produced. In the first set, the evolutionary rate was kept constant and the number of characters was varied, expecting an increase in information content with an increase in the number of characters. In the second set, the number of characters was kept constant and the evolutionary rate and hence the level of homoplasy was varied. In this set of data matrices, we would expect a decrease in information content with increasing levels of homoplasy.

Data sets were generated using Seq-Gen (Rambaut and Grassly, 1997) on a randomly chosen 17-terminal tree with equal branch lengths. An HKY85 model of sequence evolution with discrete gamma rate heterogeneity (6 categories, alpha = 0.50), ti/tv rate ratio set to 2.0, and unequal nucleotide frequencies (A: 0.30, C: 0.20, G: 0.20, T: 0.30) was applied. Tree lengths and indices for the *Drosophila* and the simulated data sets were calculated in PAUP (Swofford, 1993) excluding uninformative characters. For bootstrapping, 1000 replicates with heuristic search, TBR branch swapping, and random addition sequence (setting NREPS = 1) were used. Bremer support values were calculated using Autodecay (Eriksson and Wikström, 1996) in conjunction with PAUP (using heuristic searches with 10 random additions and TBR branch swapping).

## The Retention Index, Consistency Index and Data Decisiveness

It is clear from both the empirical and simulated data that DD does not capture information content any differently than *C* or *R* (Fig. 1 and Table 1). In fact, DD seems to reflect the same aspect of the investigated data sets as *R* (for all data sets, *R* and DD were highly correlated, *R*^2^ between 0.997 and 1.000, *p* < 0.0001). This is easily explained if there is a strong correlation between *s̅* and *G*, the number of steps on an unresolved bush (same quantity as used in calculation of *R*, Farris, 1989). Furthermore, the presence of the scaling factor in the DD formula (*s̅* – *M*; the mean homoplasy of the data set over all trees, De Laet and Smets, 1999), could explain why DD is sensitive to changes in branch length (Fig. 1a) but not to changes in the number of informative characters (Fig. 1b).

**Figure 1:**
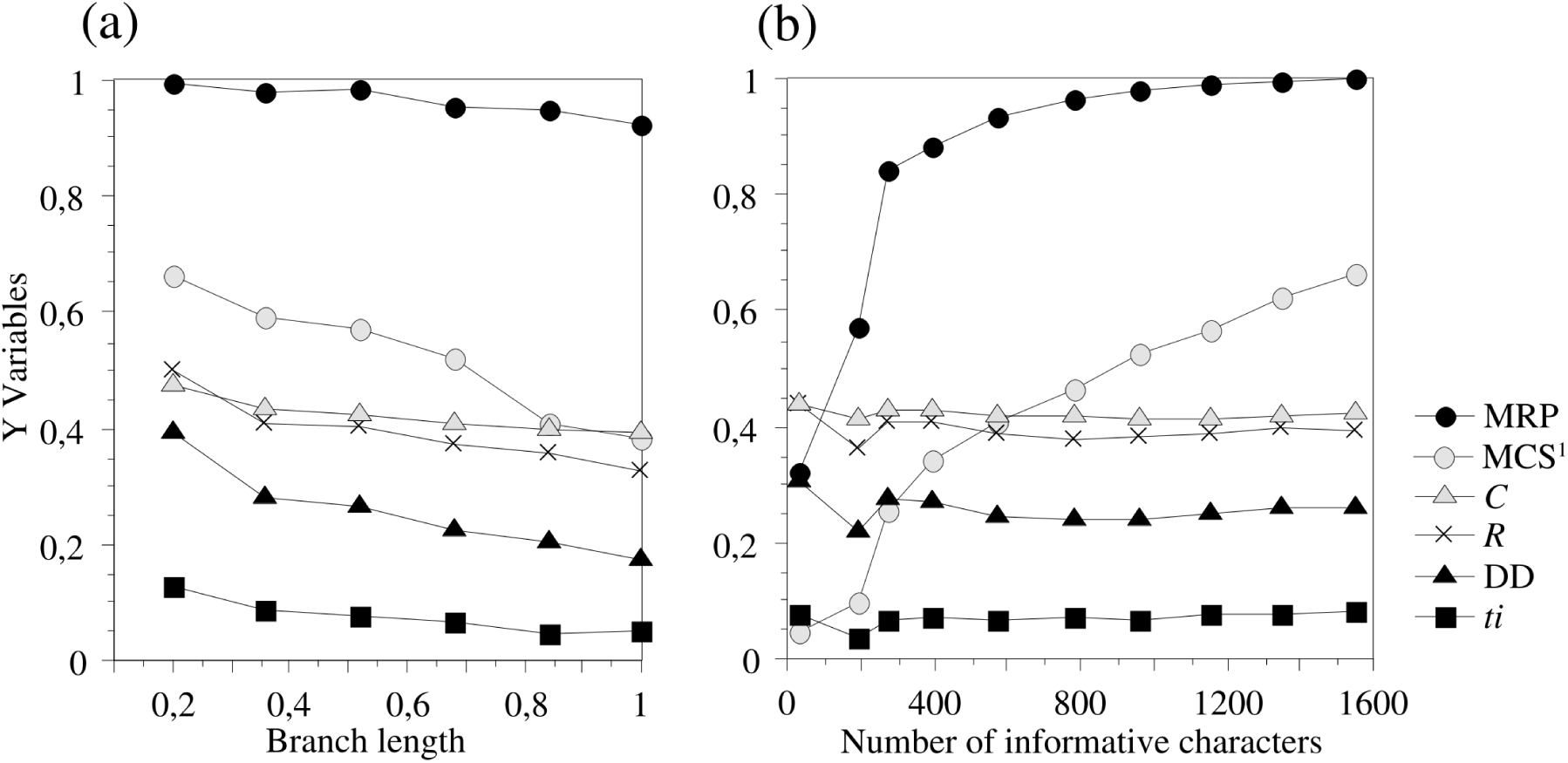
Simulated data sets illustrating various indices of data quality and their dependence on (a) the level of homoplasy (expressed as branch length or the expected number of substitutions per site for each branch), and (b) on the number of informative characters. Data sets in Fig 1b were generated as in Fig. 1a (see text for details), except that branch lengths were held constant at a value of 0.5. (^1^ Notice that mean character support, MCS, is scaled by ten to fit the graph.)

**Table 1.** Table showing results for the eight partitions of the *Drosophila* data (Baker and DeSalle, 1997) and all partitions combined.

| Data set | *n*<sub>tot</sub><sup>a</sup> | *n*<sub>info</sub><sup>b</sup> | *n*<sub>MPT</sub><sup>c</sup> | *n*<sub>nodes</sub><sup>d</sup> | Length | *C* | *R* | *ti* | DD | MRP | *s*<sub>RP</sub> | MCS | *s*<sub>CS</sub> |
|---|---|---|---|---|---|---|---|---|---|---|---|---|---|
| Combined | 2550 | 499 | 1 | 14 | 1415 | 0.499 | 0.496 | 0.177 | 0.417 | 0.906 | 0.145 | 4.153 | 2.199 |
| hb | 473 | 120 | 13 | 10 | 276 | 0.638 | 0.676 | 0.210 | 0.631 | 0.742 | 0.310 | 2.565 | 2.094 |
| 16S | 360 | 42 | 33 | 6 | 98 | 0.541 | 0.628 | 0.163 | 0.554 | 0.462 | 0.285 | 1.128 | 1.578 |
| ACHE | 323 | 60 | 4 | 12 | 154 | 0.578 | 0.536 | 0.175 | 0.470 | 0.518 | 0.291 | 1.067 | 1.025 |
| ADH | 222 | 58 | 6 | 10 | 154 | 0.565 | 0.571 | 0.130 | 0.510 | 0.498 | 0.288 | 0.947 | 0.877 |
| ND1 | 148 | 19 | 26 | 5 | 50 | 0.560 | 0.639 | 0.200 | 0.580 | 0.426 | 0.311 | 0.807 | 0.878 |
| wg | 429 | 65 | 3 | 10 | 168 | 0.524 | 0.538 | 0.083 | 0.466 | 0.440 | 0.268 | 0.750 | 0.682 |
| COIII | 229 | 58 | 5 | 7 | 197 | 0.406 | 0.448 | 0.066 | 0.356 | 0.351 | 0.259 | 0.691 | 1.121 |
| COII | 366 | 77 | 12 | 7 | 252 | 0.417 | 0.372 | 0.060 | 0.269 | 0.340 | 0.258 | 0.514 | 0.504 |

<sup>a</sup> Total number of characters.
<sup>b</sup> Number of informative characters.
<sup>c</sup> Number of most parsimonious trees (MPTs).
<sup>d</sup> Number of nodes in MPT/strict consensus.

## Measures Based on Bremer Support

Bremer support (also known as decay index or branch support) measures the degree to which clades are resolved by a data set. However, Bremer support values are not directly comparable across data sets because they must be judged in relation to the number of informative characters. A Bremer support of, say, 30 implies a different level of confidence in a 100-character data set than in a 30,000-character data set (see also Lee, 2000; Oxelman et al., 1999). The Bremer support values cannot be normalized by simply comparing them with the total number of informative characters in the matrix. This is because each value is more strongly dependent on the number of characters informative about the branching order in the local tree surrounding the node it is associated with, rather than on the total number of informative characters in the matrix. Assume, for instance, that the most parsimonious tree consists of two subclades of three taxa each, and that the interior branch in each of these subclades has a Bremer support of 5. Assume further that there are 100 characters informative about the relationships in one of the subclades and 5 about relationships in the other subclade. A Bremer support of five could easily have arisen by chance in a set of 100 characters, but not in a set of 5 characters. Clearly then, the relationships are more convincingly resolved in the subclade with fewer informative characters, even though the Bremer support and the total number of characters are the same.

Any measure of data quality derived directly from Bremer support values will suffer from these adjustment problems. For instance, this applies to total support *t*, (Källersjö et al., 1992) which is simply the sum of the Bremer support values in the shortest tree. It also applies to other statistics that could be derived from the Bremer support values of the shortest tree, such as the frequency distribution, mean and variance.

One possibility of addressing the problem is to compare the Bremer support with some value or distribution of values that we might expect to achieve by chance (Källersjö et al., 1992) but this approach is significantly more computationally complex than calculating jackknife or bootstrap values and has not been widely used. Another possibility is to scale total support by the length *s* of the most parsimonious tree(s), giving the total support index *ti* = *t*/*s* (Bremer, 1994), but this scaling destroys the correlation between the index and information content. Rather than measuring information content, the total support index reflects average character quality, like *C*, *R* and DD. This is clearly shown in the simulated sets of data matrices (Fig. 1), where *ti* follows the same trend as *C*, *R* and DD. Like the other indices, *ti* also fails to pick up the increased information content in the total evidence analysis of the *Drosophila* data (Table 1).

## A Simple Measure Based on Resampling

In contrast to Bremer support values, resampling proportions (RPs) are more easily comparable within and across data sets. Referring to the example discussed above, a clade with a Bremer support of 30 will tend to be associated with a high RP in a 100-character matrix but with a low RP in a 30,000-character matrix. This property makes RPs interesting as the basis of a measure of information content.

The simplest way of using RPs in summarizing information content is to calculate some statistic directly from the raw values. To my knowledge, the earliest attempt to do this is that of Oxelman and Eriksson (1998). They used the sum of RPs on the bootstrap 50% majority rule consensus as an indicator of data information content. However, although this descriptive statistic might be useful for individual data sets, it cannot be compared across data sets with different numbers of taxa. This suggests a measure that scales the sum of the RPs to the number of potentially supported branches in the tree. This measure will be referred to as the *Mean Resampling Proportion*, MRP, and is defined as

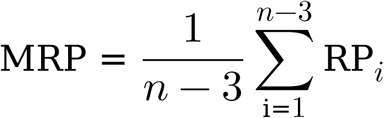

where *n* is the number of terminals and RP*_i_* is the resampling proportion of clade *i*. In calculating MRP, I suggest focusing on the tree giving the highest value of MRP. This tree can be approximated effectively by adding partitions to a 50% majority-rule consensus in order of decreasing RPs, until the tree is fully resolved. Another possibility is to use the most parsimonious tree. If there are several equally parsimonious trees, the average value over them could be used. To simplify calculations, one of the MPTs can be chosen arbitrarily and used for the purpose of estimating MRP. The variance in MRP estimates obtained from different, equally parsimonious trees should be negligible, since most of the contribution to the MRP value comes from the well-supported clades that are likely to be present in all of the most parsimonious trees (for example, MRP for the 33 MPTs from the 16S *Drosophila* data has a variance of 0.0003 and a range of 0.075).

The MRP value gives a simple key to the information content of the underlying data matrix. A matrix producing a tree with RPs of near 100 % for all groups would have MRP close to 1 whereas lower clade probabilities will tend to lower MRP. MRP measures the average level of support but in many contexts, the distribution of the support among clades, or the variance among clade probabilities, is equally important. For instance, a data set with an intermediate MRP value may either have the support for clades highly localized, in which case a few RPs are high but the rest are low resulting in significant variance around the mean. Conversely, the support may be more evenly spread throughout the tree, in which case the values of different clades are more similar and the variance is lower. The distribution of RPs in the summary tree can be illustrated in a graph that provides a powerful description of the type of information content in the data (Fig. 2). It shows how the information is partitioned into support for different clades. The distribution can be summarized by calculating the standard deviation of RP values, *s*_RP_. If desirable, this value may be scaled by the mean to obtain the relative standard deviation. Other statistics derived from the distribution may be relevant, e.g., the number of clades with an RP value above a certain threshold (e.g., Sanderson, 1989; Björklund, 1999).

**Figure 2:**
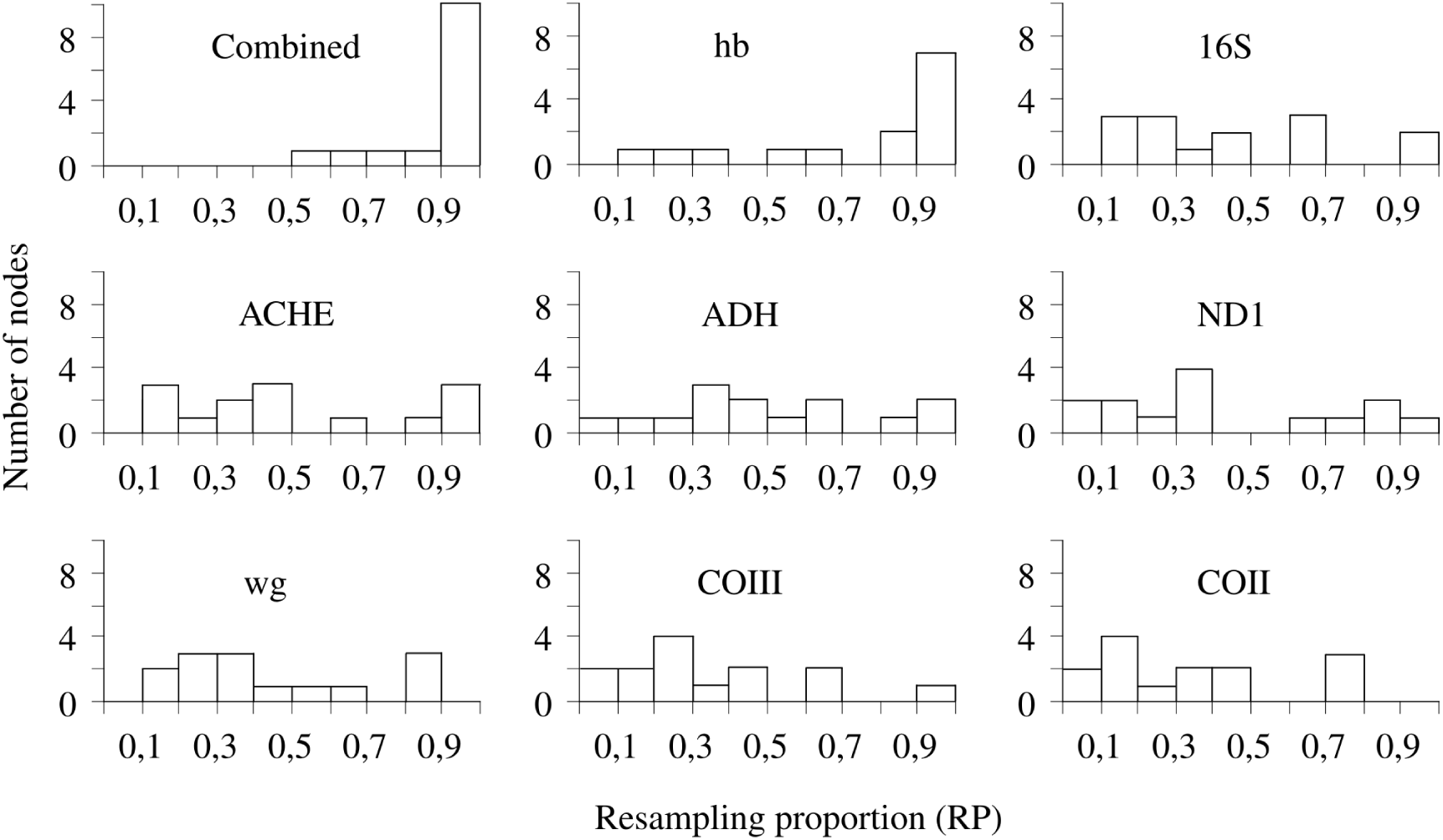
The distribution of resampling proportions, RP, as summaries of the information content in the *Drosophila* data matrices.

## An Improved Measure Based on Resampling

As expected, MRP more accurately reflects the information content in data matrices than the other measures discussed so far. For instance, it clearly signals the increased information content in the total-evidence analysis of the *Drosophila* data sets; the MRP values for single data sets vary between 0.340 and 0.742 whereas the combined data set has a MRP of 0.906 (Table 1). Note that there is a weak correlation between MRP on one hand and the other indices on the other for the single data sets. However, when the indices differ, it is the MRP values that more accurately reflect the degree to which the data sets resolve relationships with high confidence. For instance, the ACHE data set has a comparable *C* and a lower *R* (and DD) than the ND1 data set and yet, the former resolves relationships better. This is indicated among other things by the smaller number of most parsimonious trees for the ACHE data set (4 versus 26). This difference is accurately reflected by MRP, which is higher for the ACHE than for the ND1 data set (0.518 versus 0.426).

The simulated data sets also illustrate that MRP is correlated with information content. MRP values decrease with increasing levels of homoplasy (Fig. 1a) but increase with the number of informative characters (Fig. 1b). The latter curve, however, also illustrates a problem with MRP. Note that the MRP values increase rapidly when we go from small to intermediate data sets (50 to 250 informative characters) but that the increase slows down considerably as the data sets grows larger and the MRP values get beyond 0.80. This occurs because support is not linearly related to clade probabilities. There is a considerable difference in support between clades having probabilities of e.g. 0.90 and 0.95, whereas probabilities of 0.60 and 0.65 are associated with support levels that can hardly be distinguished. This also means that the arithmetic mean of the RPs underestimates the average support level, since the RPs do not influence MRP to the extent that they should.

A standard solution to a problem of this kind is rescaling or transformation of the raw measurements. The fact that it is desirable to stretch out the scale close to 1 suggests that a suitable transformation of the raw RPs, is provided by ln (1 / (1 – RP)). This transformation has an interesting property. For standard resampling schemes, including the bootstrap and jackknife, it approximates the number of characters that would be expected to give the observed support value in a homoplasy-free binary matrix analyzed with parsimony. In this situation, a clade will have an expected bootstrap proportion of RP = 1 – (1 – *r* / *n*)*^n^*, where *r* is the number of characters supporting the clade and *n* is the total number of characters in the data set (Felsenstein, 1985). When *n* becomes large, this value approaches 1 – *e*^−^*^r^* (Harshman, 1994) and thus *r* = ln (1 / (1 – RP)) (Fig. 3). The expected jackknife proportion is 1 – *p^r^*, where *p* is the probability of a character being absent from any one pseudoreplicate. By setting *p* = *e*^−1^, the expected jackknife proportion will be the same as the proportion approached by the bootstrap in large data sets (Farris et al., 1996).

**Figure 3:**
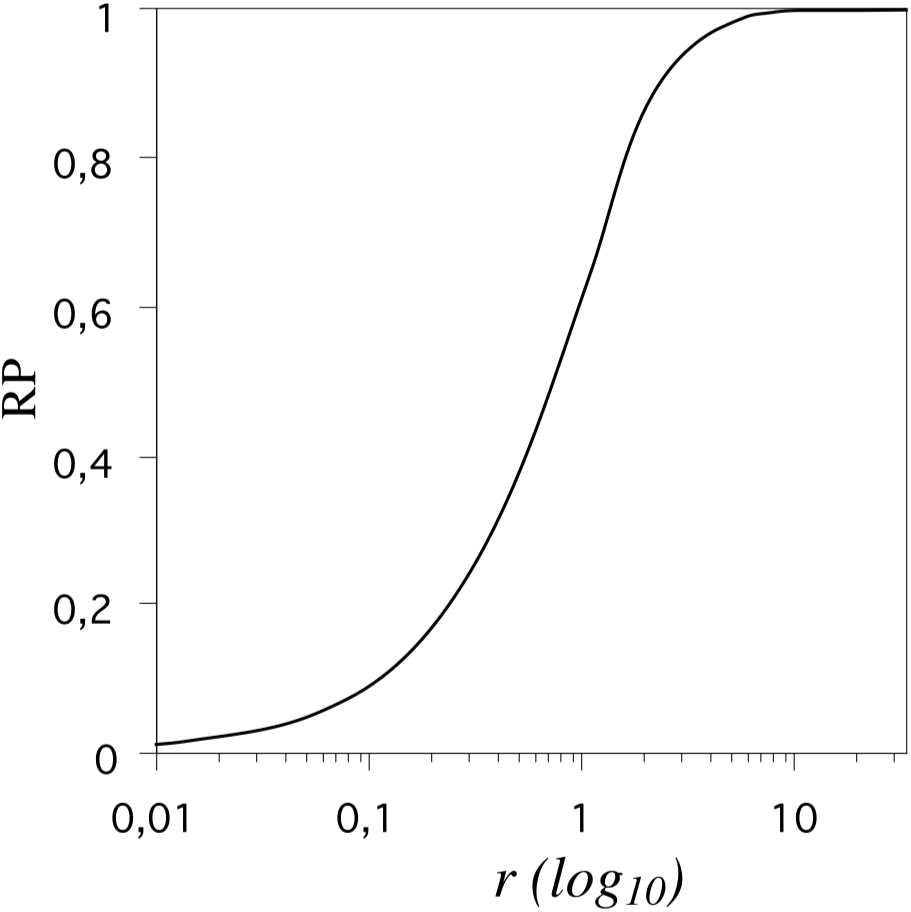
The relationship between the number of characters supporting a clade, *r*, and the resampling proportion, RP, for that clade in a data matrix free of homoplasy. The graph is based on the approximate formula RP = 1 – *e*^−^*^r^*. Note that it takes proportionally more characters for increasing the RP for a clade having e.g. RP = 0.90, compared to one having RP = 0.50.

Since it can be interpreted as the number of unambiguous characters supporting a clade, the ln (1 / (1 – RP)) transformation provides a powerful yet intuitive measure of clade support. The same transformation can be used to obtain an efficient matrix representation of the original data (Ronquist, 1996). Assume that the tree summarizing clade support values is represented as a matrix of binary characters, each supporting a unique partition in the tree, and that each character is weighted according to the transformed RPs (ln (1 / (1 – RP)). Under parsimony, we can evaluate the length of any tree using the weighted representation matrix and be sure to get a value that is highly correlated with the length of the same tree using the original data (Ronquist, 1996). The ln (1 / (1 – RP)) transformation also describes how RPs add up in combined analyses of different data sets (at least in the absence of character conflict). For example, if two different data sets both give a RP of 0.75 for a particular clade, the RP for the clade in the combined analysis will be 0.94 since ln (1 / (1 – 0.94)) = ln (1 / (1 – 0.75)) + ln (1 / (1 – 0.75)). Thus, weighted representation matrices provide a highly precise way of combining separate analyses into supertrees, far more sophisticated than simply combining unweighted matrices describing the topology of the original trees.

Because RPs are estimated from a finite number of replications, some clades may occur in the shortest trees in all pseudoreplicates. This makes it impossible to calculate the corresponding number of supporting characters since ln (1 / (1 – RP)) is undefined when RP = 1. The heart of the problem is that there are no trees in the sample violating the monophyly of the clade although at least some incompatible trees are expected if the sample is large enough. A standard solution to such problems, where the maximum likelihood estimate of zero probability is inappropriate, is to use a Bayesian estimate where pseudocounts are added to describe a prior expectation of observing at least one data point of each type (Durbin et al., 1998) Assume that we observe *n* trees having a particular clade in our resampling tree set of *N* trees. The Bayesian estimate of the RP would then be RP = *n* / (*N* + 1). This means a small correction downwards of RP values close to 1 but a negligible correction for RP values close to 0. For illustration, suppose that a clade occurred in 100 of 100 replicates in a bootstrap analysis. The estimated RP would then be 100 / 101 = 0.99. If the clade occurred in 1,000 of 1,000 replicates, the estimate would be 1,000 / 1,001 = 0.999.

The distribution of transformed RPs can be illustrated in a graph (see Fig. 5 for the *Drosophila* data) or summarized by descriptive statistics, such as the mean and standard deviation. The mean will be referred to here as the *Mean Character Support* (MCS) and is estimated as

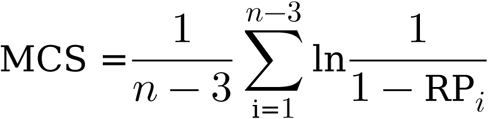

The corresponding standard deviations will be referred to as the *standard deviation of character support* (*s*_CS_). As for raw RPs, the distribution of character support in the summary tree can be illustrated in a graph (Fig. 4).

**Figure 4:**
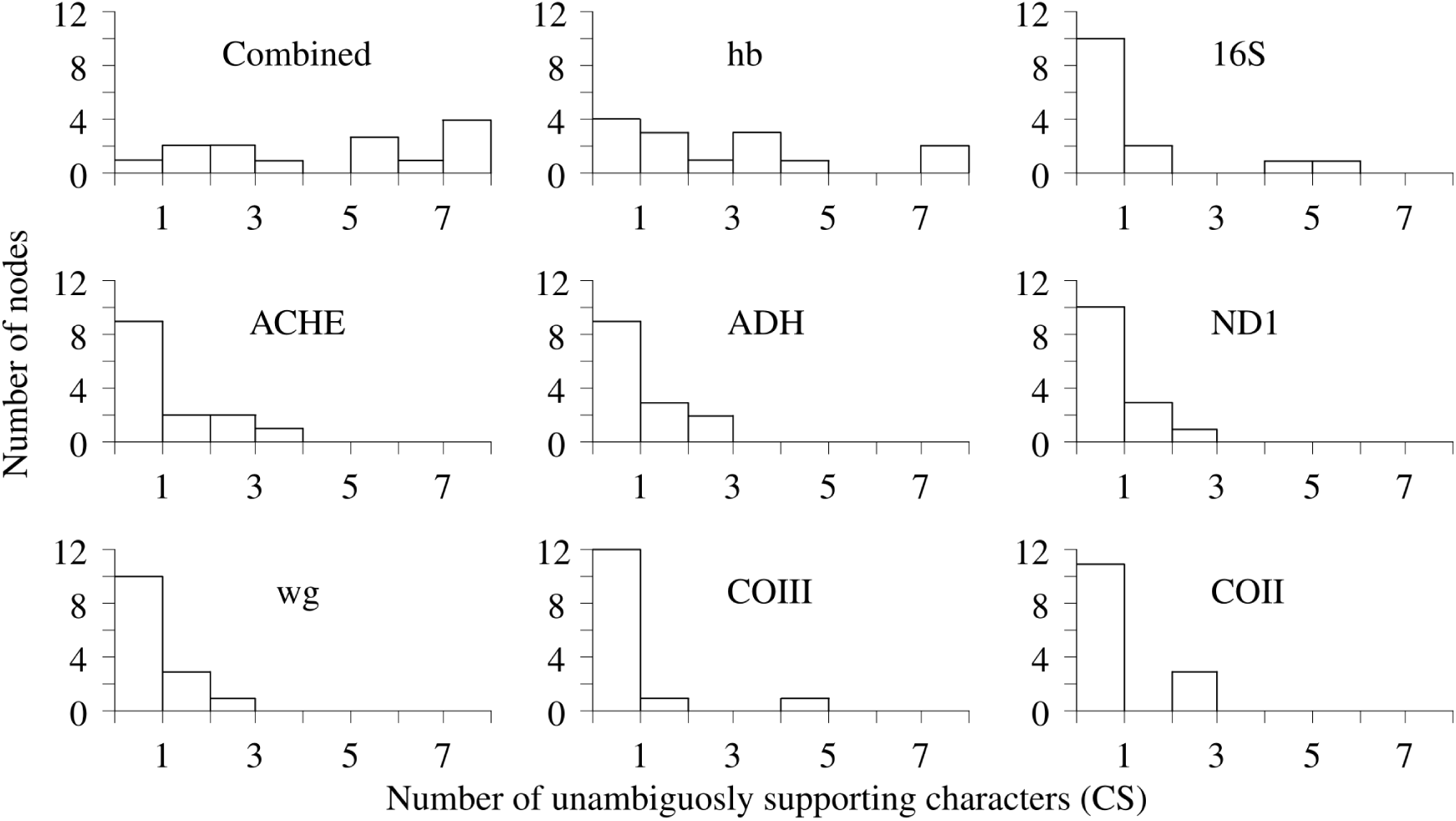
The distribution of character support, CS, as summaries of the information content of the *Drosophila* data matrices.

MCS is based on the approximate formula for describing the relationship between *r* and RP. Unless the total size of the representation matrix is very small, this approximation is quite accurate. For instance, assume that there are 17 taxa in the data set and that the approximate formula gives a support of 0.50 characters (RP = 0.39) for each of the 14 interior nodes (approximately corresponding to the smallest representation matrix analyzed here, containing roughly 7 characters in total; COII, Table 1). Application of the exact formula shows that the true support in this case is 0.48 and not 0.50 characters per internal node, a deviation of only 4%.

MCS provides a more sensitive measurement of information content than MRP. For instance, the partitions of the *Drosophila* data have MCS values ranging from 0.514 to 2.565, while the combined data set has an MCS value of 4.153. This large increase in information content in the combined analysis is not picked up by the MRP because this measure is underestimating the information content in really large or very good data sets. This phenomenon is evident in the sets of simulated data matrices. MCS increases linearly with the number of informative characters whereas MRP soon reaches an asymptote (Fig. 1b). Both MCS and MRP are sensitive to the level of homoplasy but the MRP values become less sensitive as the level of homoplasy decreases and the support values approach 1, whereas MCS continues to signal differences in information content even between the best data sets.

Simple summary measures like MCS and *s*_CS_ cannot reflect all qualities of the underlying distribution. For instance, a data set with one highly and many poorly supported clades may have the same MCS and *s*_CS_ as a data set with varied, but mostly intermediate RPs. The difference between the data should, nevertheless, be clearly visible when comparing the underlying distribution of clade support values. Although this example might be seen as a shortcoming of the MCS, it merely elucidates the fact that “data quality” is a complex quantity to measure, and no single index can capture all aspects of it.

## Conclusions

Mean character support (MCS) is an appropriate measure of the degree to which data lend support to hypotheses about the monophyly of clades. It effectively summarizes two of the most important aspects of a phylogenetic result: resolution and support. It is easily calculated from resampling proportions, once these have been estimated. MCS has an easy and intuitive interpretation: it represents the mean number of characters supporting clades in a binary representation matrix of the original data set. Unlike other measures of data quality, MCS is easily applicable to different phylogenetic inference methods, such as distance, parsimony, and maximum likelihood approaches.

## Software Availability

Software for calculating MCS, MRP, and their standard deviations is downloadable from https://github.com/nylander/get_newick_info.

## Acknowledgements

This research was supported by a grant from the Swedish Natural Science Research Council (621-2001-2963) to Fredrik Ronquist.

## Notes

https://github.com/nylander/get_newick_info

### Competing Interest Statement

The authors have declared no competing interest.

